# Switching between newly learned motor skills

**DOI:** 10.1101/2024.03.22.586357

**Authors:** Kahori Kita, Yue Du, Adrian M. Haith

## Abstract

Studies of cognitive flexibility suggest that switching between different tasks can entail a transient switch cost. Here, we asked whether analogous switch costs exist in the context of switching between different motor skills. We tested whether participants could switch between a newly learned skill associated with a novel visuomotor mapping, and an existing skill associated with an intuitive mapping. Participants showed increased errors in trials immediately following a switch between mappings. These errors were attributable to persisting with the pre-switch policy, rather than imperfect implementation or retrieval of the post-switch policy. A subset of our participants further learned a second new skill. Switching between these two novel skills was initially very challenging, but improved with further training. Our findings suggest that switching between newly learned motor skills can be challenging, and that errors in the context of switching between skills are primarily attributable to perseveration with the wrong control policy.

## Introduction

In everyday life, we frequently need to switch between different skills. For instance, when picking up and using different tools, when getting on or off a bike, or when driving a car forwards versus in reverse. In cognitive science, it is widely recognized that switching between different tasks entails a transient cost in the form of temporarily slowed performance immediately after the switch. Here, we examined whether similar switch costs apply in the context of switching between different learned motor skills and, further, what underlying processes might account for any switch costs that do exist.

The phenomenon of task switching has been extensively studied in cognitive science (Rogers and Monsell, 1995; Gopher, 1996; Meiran, 1996; Vandierendonck et al., 2010; Braem et al., 2019; Hazeltine, 2024). In a typical experiment, participants might learn to perform two or more different tasks that require them to respond to the same stimuli according to different rules. For example, participants might be asked to switch between classifying a digit as odd or even and classifying it as high or low (Sudevan and Taylor, 1987; Rogers and Monsell, 1995; Logan and Bundesen, 2003). Though each individual task is simple, participants generally exhibit longer reaction times and/or are more prone to errors immediately after switching between tasks. Two different explanations for such switch costs have emerged (Monsell, 2003, 2017; Vandierendonck et al., 2010). One set of theories has suggested that switch costs arise from the need to retrieve a new task rule or policy from long-term memory after a switch (Mayr and Kliegl, 2000; Rubinstein et al., 2001; Sohn and Anderson, 2001; Schneider and Logan, 2009) and/or performing necessary reconfiguration of neural circuits to implement the new policy (Rogers and Monsell, 1995; Logan and Gordon, 2001; Monsell and Mizon, 2006). An alternative view, however, posits that switch costs may primarily arise from difficulty in suppressing the existing task policy (Allport et al., 1994; Wylie and Allport, 2000; Monsell, 2003; Kiesel et al., 2010). Either or both of these explanations might apply in the context of switching between motor skills.

The need to switch between motor skills should be equally important as for cognitive skills, but has not been examined as closely. As described above, task switching in the cognitive domain is often studied by switching task rules that require different responses to the same stimuli. A direct analog of this in the context of motor control is to switch between different visuomotor mappings, requiring different motor commands in a given state of the task (e.g. to reach a given target location). Along these lines, numerous studies have examined the ability to switch between two opposing perturbations, such as perceptual distortions that rotate the visual field clockwise versus counter-clockwise (Tong et al., 2002; Osu et al., 2004; Krakauer et al., 2005; Addou et al., 2011; Forano et al., 2021). In these cases, however, people are generally unable to switch between these perturbations at all and instead need to re-learn the newly imposed perturbation each time (though see (Cunningham and Welch, 1994; Huberdeau et al., 2019)). This inability to switch appears to be inherent to the implicit adaptation mechanism which is responsible for countering the perturbation (Huberdeau et al., 2015; Morehead and Xivry, 2021). It is increasingly appreciated, however, that adaptation is not the only, or even the main, learning mechanism involved in learning new motor skills. Most motor skills cannot be acquired simply by recalibrating an existing controller and instead appear to be learned by creating a new controller – a process referred to as “*de novo”* learning. Critically, unlike adaptation, *de novo* learning leads to minimal aftereffects (Yang et al., 2021; Haith et al., 2022; Gastrock et al., 2023). Therefore, although participants cannot switch between different motor behaviors learned through adaptation, they may be able to switch between behaviors learned through *de novo* learning.

Here, we employ a motor learning task which has been shown to be learned through *de novo* learning rather than adaptation (Haith et al., 2022) how people are able to switch between different newly learned motor skills. In this task, participants use both hands to control the position of an on-screen cursor through a highly non-intuitive mapping (Yang et al., 2021; Haith et al., 2022). We asked whether participants could switch between this newly learned skill and a pre-existing baseline behavior. Subsequently, we also asked whether participants could learn multiple such skills associated with different mappings, and switch between them.

## Results

### Participants learned the De Novo mapping over multiple days of practice

Twenty-five healthy participants practiced maneuvering an on-screen cursor to reach to a series of visual targets using a novel and non-intuitive mapping between their hands and the cursor (De Novo mapping, Fig. 1A-D). As in our previous studies (Yang et al., 2021; Haith et al., 2022), participants initially found this task very challenging, but their performance gradually improved with practice across multiple sessions. Both the reaction time (reaction time of the first 5 blocks on Day 1 (mean ± s.e.m): 570.55 ± 34.87 ms, the last 5 blocks on Day 4: 289.14 ± 7.58 ms, paired t-test, P < 10^-6^, t(24) = 8.57, not shown as a figure) and the initial direction error of the cursor movement (the first 5 blocks on Day 1: 51.63 ± 1.74 °, the last 5 blocks on Day 4: 21.73 ± 1.22 °, paired t-test, P < 10^-6^, t(24) = 18.73, Fig. 1F) significantly decreased between the beginning and end of practice. By the end of Day 4, performance under the De Novo mapping was almost as good as performance under an intuitive mapping in which the cursor appeared at the average location of the two hands (“Baseline Mapping”), though reaction times remained slightly longer (reaction time under the Baseline mapping: 275.09 ± 6.98 ms, paired t-test, P = 0.0128, t(24) = - 2.69) and initial direction error was slightly greater (initial direction error under the Baseline mapping: 8.42 ± 0.56 °, paired t-test, P < 10^-5^, t(24) = -10.57). By the end of learning, participants had developed a novel movement policy associating the direction of the target to movement of the right and left hands (see example participant in Fig. 1G).

**Figure 1.**
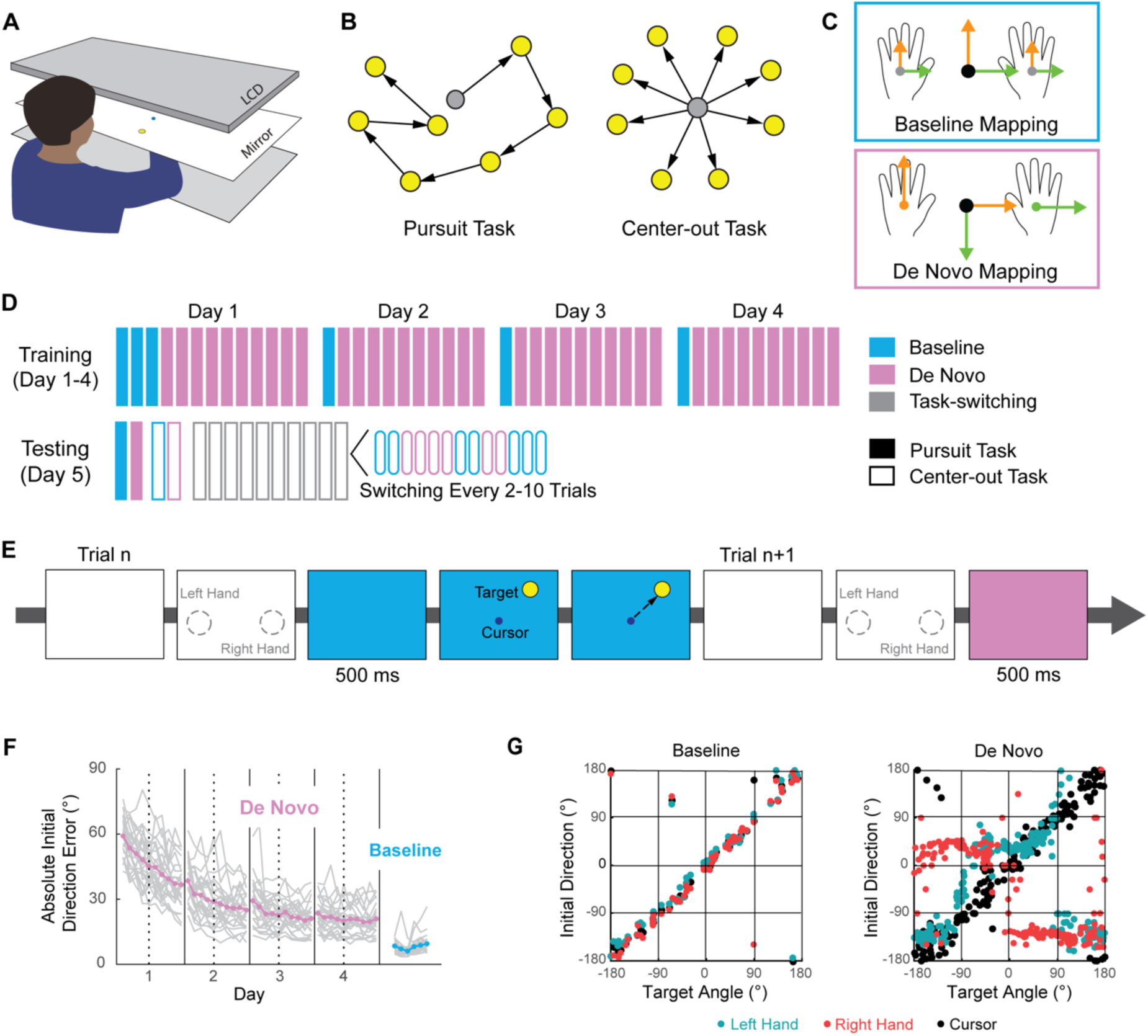
Experiment 1, switching between the De Novo and the Baseline mapping. A) participants performed reaching movements to move a cursor toward targets presented via mirrored display. B) Participants were trained on the De Novo mapping by making a series of point-to-point movements around the workspace (pursuit task). Participants’ ability to switching between mappings was assessed in a center-out task. C) Two mappings: the Baseline mapping, in which the cursor appeared at the average location of the two hands, and the De Novo mapping, in which each hand controls one degree of freedom of the cursor but orthogonal to its own movement. D) Participants first performed the Baseline mapping to measure their baseline performance, followed by four days of practice on the De Novo mapping (600 trials per day). On the fifth day, participants were tested their ability to switch between the Baseline and De Novo mappings on a trial-by-trial basis based on a color. E) During the task-switching assessment, the background color of the screen turned either blue or magenta to indicate which mapping should be used in the upcoming trial. The target appeared on the screen 500 ms later at which point participants had to rapidly initiate a movement of the cursor towards it. F) Performance during initial learning of the De Novo mapping. Thick lines indicate average across participants (magenta: De Novo mapping, blue: Baseline mapping). Thin gray lines indicate individual participants. G) Behavior of a representative participant at the center-out task on Day 5. Initial direction of the left hand (blue gray dots) and right hand (red dots) using either the Baseline or De Novo mapping, showing that the participant adopted a very different coordination pattern for the De Novo mapping compared to their behavior in the Baseline mapping. Black dots indicate initial direction of the cursor.

### Participants could easily switch between the Baseline and De Novo mappings

On the fifth day, we examined participants’ ability to switch between the learned De Novo mapping and the Baseline mapping (Testing, Fig. 1D). We used center-out reaches from a single start position so that the mapping could easily be switched between trials. We first measured participants’ performance in the center-out task when the mapping was fixed throughout a 60-trial block and we used this to establish a steady-state level of performance under each mapping. We then tested participants’ ability to switch between mappings within a block. The background color of the screen at the start of each trial indicated which mapping would be applied in that trial and this switched randomly every 2-10 trials.

Figure 2A shows data from a representative participant (Participant 1). This participant showed an increased directional error in trials immediately following a switch from the Baseline to the De Novo mapping (steady state of the De Novo mapping (mean ± std): 16.29 ± 18.78 °; the first trials post switch: 29.74 ± 23.07 °; two-sample t-test, P < 0.001, t(226) = 4.22) as well as following switches from the De Novo mapping to the Baseline mapping (steady state of the Baseline mapping: 9.43 ± 22.28 °; the first trials post switch: 24.85 ± 22.91 °; two-sample t-test, P < 0.001, t(107) = 3.55). Within 2-3 trials, however, their directional errors returned to the same steady-state level as that seen in blocks with no task switching at all (horizontal blue line). Reaction times – the measure more conventionally used to establish switch costs in cognitive tasks, did not exhibit this transient switch costs (Fig. 2A, inserted panel), likely because we deliberately pressured participant’s reaction times in order to focus on the pattern of directional errors following a task switch. These transient but consistent increases in directional error following a switch suggest a cost associated with switching between mappings, analogous to the task switch costs identified in more cognitive tasks.

**Figure 2.**
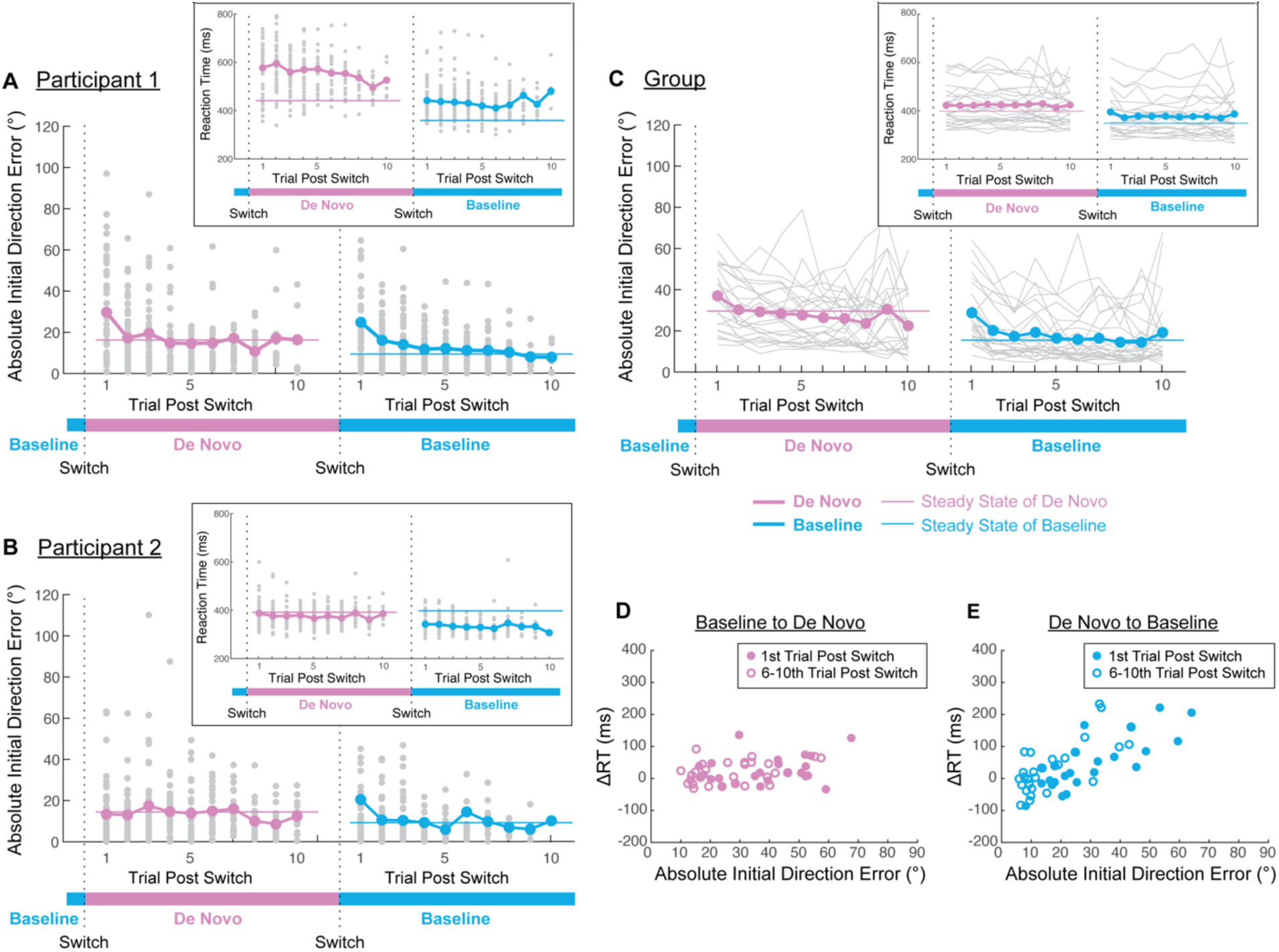
Performance when switching between the Baseline and De Novo mappings. Panels A-C show performance (absolute initial direction error and reaction time) in the first ten trials post switch either from the Baseline to the De Novo mapping or from the De Novo to the Baseline mapping, during the switching assessment on Day 5. A,B) Performance of two representative participants (A, Participant 1; B, Participant 2). Thick lines indicate average initial direction error for each participant, and thin lines indicate steady state performance for each participant, assessed in blocks without any task switching at the start of Day 5 (magenta: De Novo mapping, blue: Baseline mapping). Gray dots indicate performance in individual trials. C) Performance across all participants. Thick colored lines indicate mean performance across all participants. Thin gray lines indicate averaged behavior across trials for each individual participant. Thin magenta and light blue lines indicate steady state performance of under the De Novo and the Baseline mappings on Day 5. D,E) Relationship between absolute initial direction error and reaction time in post-switch trials when switch from the Baseline to the De Novo (D) and from the De Novo to the Baseline mapping (E). ΔRT indicates different between rection time post switch trials and steady-state performance under that mapping in nonswitching blocks (thin horizontal lines in A-C). Filled circles indicate first trials post switch and open circles represent the average across the 6-10th trials post switch. Each circle indicates one participant.

Not all participants exhibited a switch cost when the mapping changed. Figure 2B shows performance of another participant (Participant 2) who showed no increase in directional errors following switches from the Baseline mapping to the De Novo mapping (steady state of the De Novo mapping (mean ± std): 14.48 ± 14.45 °; the first trials post switch: 13.36 ± 14.48 °; two-sample t-test, P = 0.6469, t(228) = -0.46), and thus it was possible to switch immediately form the Baseline mapping to the De Novo mapping based on the color cue. Furthermore, Participant 2 had negligible changes in reaction time in the first trial after a switch either way (Fig. 2B, inserted panel).

At an overall group level, however, we found clear increased directional errors in trials immediately following a switch, both when switching from the Baseline to the De Novo mapping and vice versa (Fig. 2C, steady state for the De Novo (mean ± s.e.m): 29.60 ± 2.59 °, trials following switch from the Baseline to De Novo: 37.25 ± 3.12 °, paired t-test, P = 0.0007, t(24) = 3.91; steady state for the Baseline: 15.48 ± 3.26 °, the first trials following switch from the De Novo to Baseline: 28.97 ± 3.08 °, paired t-test, P = 0.0080, t(24) = 2.89). For trials between two and four trials following a switch, there were no significant increases in initial direction error over steady-state performance, suggesting that the switch cost was short-lived. Participants at a group level also showed no transient increases in reaction time post-switch in either direction (Fig. 2C inserted panel, from Baseline to De Novo, first trials post switch: 423.39 ± 16.72 ms; 6th-10th trials: 424.81 ± 16.95 ms; from De Novo to Baseline, first trials post switch: 395.999 ± 17.39 ms; 6th-10th trials: 383.430 ± 18.01 ms).

Although we did not observe transient increases in reaction time following a switch, we did find that many participants did exhibit a small but consistent increase in reaction time throughout the task-switching assessment, compared to their reaction times in blocks with the same mapping but without any task switching (Fig. 2C, inserted panels). We wondered whether this increase in reaction times might be related to task errors, perhaps if increasing reaction times enabled participants to reduce their directional errors. We examined the relationship between the magnitude of these elevated reaction times and increases in directional error of the cursor, potentially to see if participants who avoided making errors did so by having overall increased reaction times. However, we found that this was not the case and in fact found the opposite relationship (Fig. 2DE). During switches from Baseline to De Novo mapping, there was no correlation between increases in reaction time over steady-state and increases in initial direction error either in trials immediately following a switch of mapping (Fig. 2D, correlation coefficient r = 0.20, P < 0.001). When switching from the De Novo to Baseline mapping, increases in reaction time were actually positively correlated with initial direction error (r = 0.78, P < 0.001), such that participants who increased their reaction times during the task switching blocks actually performed worse (Fig. 2E). Both of these patterns were similar 6-10 trials after a switch (Fig 2DE, open circles. From Baseline to De Novo, r = 0.17, P < 0.001; from De Novo to Baseline, r = 0.58, P < 0.001), suggesting that the overall increases in reaction time were unrelated to transient increases in directional error following a switch.

### Directional errors following a switch arose from inappropriately persisting with the pre-switch policy

Overall, our data demonstrate that most participants had little difficulty switching between the De Novo and Baseline mappings. Nevertheless, most participants did exhibit temporary increased directional errors in trials immediately following a switch. We reasoned that these errors could have been due either to imperfect execution of the new mapping – perhaps because the new control policy had not yet been fully retrieved from long-term memory and prepared for use (Monsell, 2003) – or they might have been due to participants inappropriately persisting with the policy that had been appropriate for the pre-switch mapping.

In order to dissociate between these two explanations, we closely examined the direction of movement of participants’ left and right hands, rather than just the direction of movement of the cursor. The patterns of movement of the two hands exhibited a distinctive pattern under each mapping (Fig. 1G), and we exploited this structure to infer which policy (Baseline or De Novo) participants were applying in each trial, or whether their movements were inconsistent with either policy (Fig. 3A). We used a maximum likelihood approach to estimate, on a trial-by-trial basis, the probability that participant used each policy. Figure 3C-F shows the individual hand movement directions and inferred choice of policy for the two representative participants from Figure 2, when switching from the Baseline mapping to the De Novo mapping. Figure 3GH show the choice of policy across participants when switching in both directions. In both cases, the probability of an execution error (in which behavior did not match either the Baseline or the De Novo policy) did not vary with increasing numbers of trials post-switch. Therefore, the brief increase in directional errors following a switch (Fig. 2C) was mostly driven by a small probability of incorrectly persisting with the policy for the pre-switch mapping, rather than any difficulty in retrieving and successfully implementing the post-switch mapping.

**Figure 3.**
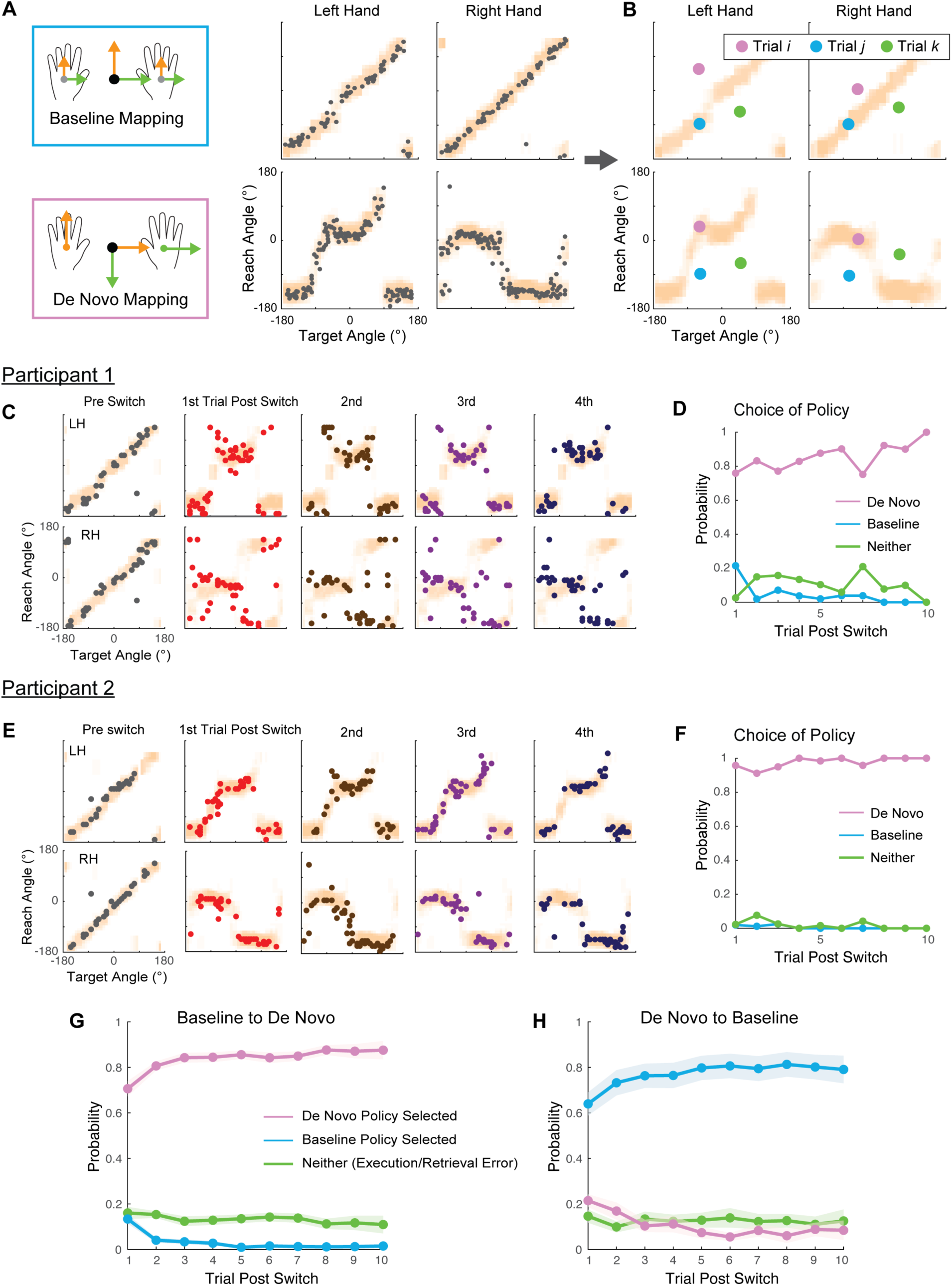
Inferring the policy used by participants on each trial post switching. A) Initial direction of the left hand and right hand varied systematically as a function of target direction (gray dots; each dot = 1 trial, taken from non-switching block at the start of Day 5). Based on this data, we estimated the conditional distribution of each hand’s direction given the target direction (orange shaded region). We did this separately for the left and right hand (left/right column) and for the Baseline and the De Novo mapping (top/bottom row). B) We used these distributions to infer, on subsequent trials, which policy participants employed. The colored circles represent potential observations (initial directions of the left and right hands given the target direction), with each trial (*i*, *j*, *k*) shown in a different color. Trial *i* (magenta) falls within the distribution of movements under the De Novo mapping (bottom row), but not the Baseline mapping (top row). We therefore infer a high likelihood that this trial was generated using the policy learned for the De Novo mapping. Trial *j* (blue) falls within the distribution of movements under the Baseline mapping (top row), but not the De Novo mapping (bottom row), so we infer a high likelihood that this trial was generated by the policy used for the Baseline mapping. Trial *k* (green) does not fall within either distribution, and we infer that this movement was not likely to have been generated under either policy. C-F) Illustration of this analysis for the two representative participants in Figure 2. (C, E) Each column shows behavior in different trials relative to a switch from the Baseline to the De Novo mapping. Each row shows a different hand (LH: left hand, RH: right hand). (D, F) shows the inferred probability of each mapping (or neither mapping) as a function of trials post switch. For Participant 1 (C, D), their behavior after the switch was reasonably well aligned with the De Novo policy in most trials. However, in a subset of trials immediately following a switch, their hand movements were more consistent with the Baseline policy. Participant 2 (E, F), by contrast, successfully applied the appropriate policy with high probability, even from the first trial. G,H) Average probability of applying each policy across participants when switching from the Baseline to the De Novo mapping (G) and from the De Novo to the Baseline mapping (H). (magenta = De Novo policy, blue = Baseline policy, green = neither policy). Shaded regions indicate ± s.e.m.

### Participants could learn a second De Novo mapping without interference

In the first experiment, we found that people had little difficulty in switching between a newly learned skill (the De Novo mapping) and a well-established skill (the Baseline mapping). However, we suspected that it may be more challenging to switch between two newly learned skills. We therefore conducted a further experiment to test whether people could learn two De Novo mappings and switch between them in a similar manner to switching between the De Novo and the Baseline Mappings.

Seven participants who had already learned the De Novo mapping in Experiment 1 (which we will now refer to as Mapping A) went on to participate in Experiment 2, in which they learned another De Novo mapping (Mapping B). Mapping B was similar to Mapping A, except the roles of the left and right hand were transposed (Fig. 4). Participants practiced Mapping B over five days (Fig. 4, Days 6-9, switching test between the Baseline mapping and Mapping B on Day 10). Learning of Mapping B proceeded in a similar manner as initial learning of Mapping A in terms of initial direction error (Fig. 5AB, the last five blocks of practice Day 4 under Mapping A (mean ± s.e.m): 19.49 ± 1.90 °, the last five blocks of practice Day 9 under Mapping B: 17.32 ± 1.36 °, paired t-test, P = 0.2169, t(6) = 1.38) and reaction time (Mapping A (mean ± s.e.m): 295.40 ± 18.43 ms, Mapping B: 325.43 ± 22.93 ms, paired t-test, P = 0.0723, t(6) = -2.18). Thus, people were able to learn a second De Novo mapping, and there did not appear to be any interference, positive or negative, from prior learning of Mapping A on subsequent learning of Mapping B.

**Figure 4.**
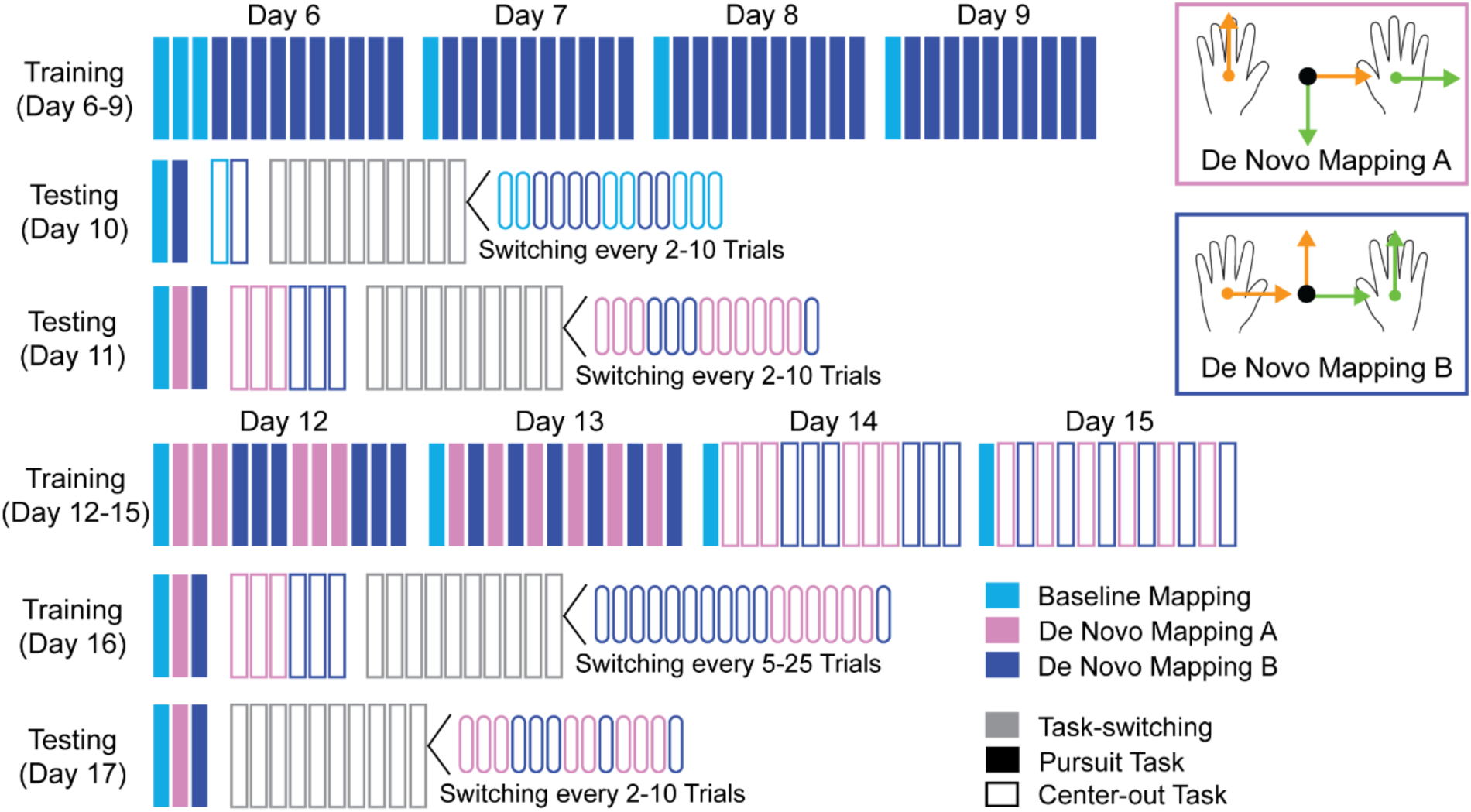
Experiment 2 – switching between two De Novo mappings. After the end of Experiment 1 (Days 1-5), a subset of participants learned a new De Novo mapping (Mapping B) (600 trials per day, Days 6-9). On Day 10, we tested participants’ ability to switch between the Baseline mapping and De Novo mapping B on a trial-by-trial basis based on a color cue. On Day 11, we assessed participants’ ability to switch between the two De Novo mappings. From Day 12 to Day 16, participants practiced switched between mappings. On Day 12, the mapping switched every three blocks with participants performing the pursuit task (Fig. 1B). On Day 13, the mapping switched every block. On Days 14-15 we repeated this process with center-out task rather than pursuit task (Fig. 1B). On Day 16, the mapping switched every 5-25 trials with center-out task. Finally, on Day 17, we repeated the assessment with the mapping switching every 2-10 trials. Participants performed 6 blocks of 60 trials for each mapping in total on Days 12-15. Two of seven participants did not complete the initial switching assessment on Day 11 and instead directly started switching training (Days 12 – 16).

On Day 11, we checked whether participants still retained the ability to perform under Mapping A after having learned Mapping B. Participants performed three blocks of the center-out task under Mapping A, followed by three blocks under Mapping B (Fig. 4, Day 11) (NB – this analysis included five participants since two participants did not complete a switching assessment on Day 11). The reaction time and initial direction errors (Fig. 5C) were similar for both mappings (Mapping A reaction time (mean ± s.e.m): 481.57 ± 54.68 ms, Mapping B reaction time: 455.28 ± 51.10 ms, paired t-test, P = 0.1630, t(4) = 1.71; Mapping A initial directional error: 35.80 ± 6.63 °, Mapping B initial directional error: 26.98 ± 2.49 °, P = 0.1599, t(4) = 1.72), and participants maintained distinct hand movements for each mapping. Therefore, participants were still able to control the cursor under Mapping A, and there was no retrograde interference from intervening learning of Mapping B.

**Figure 5.**
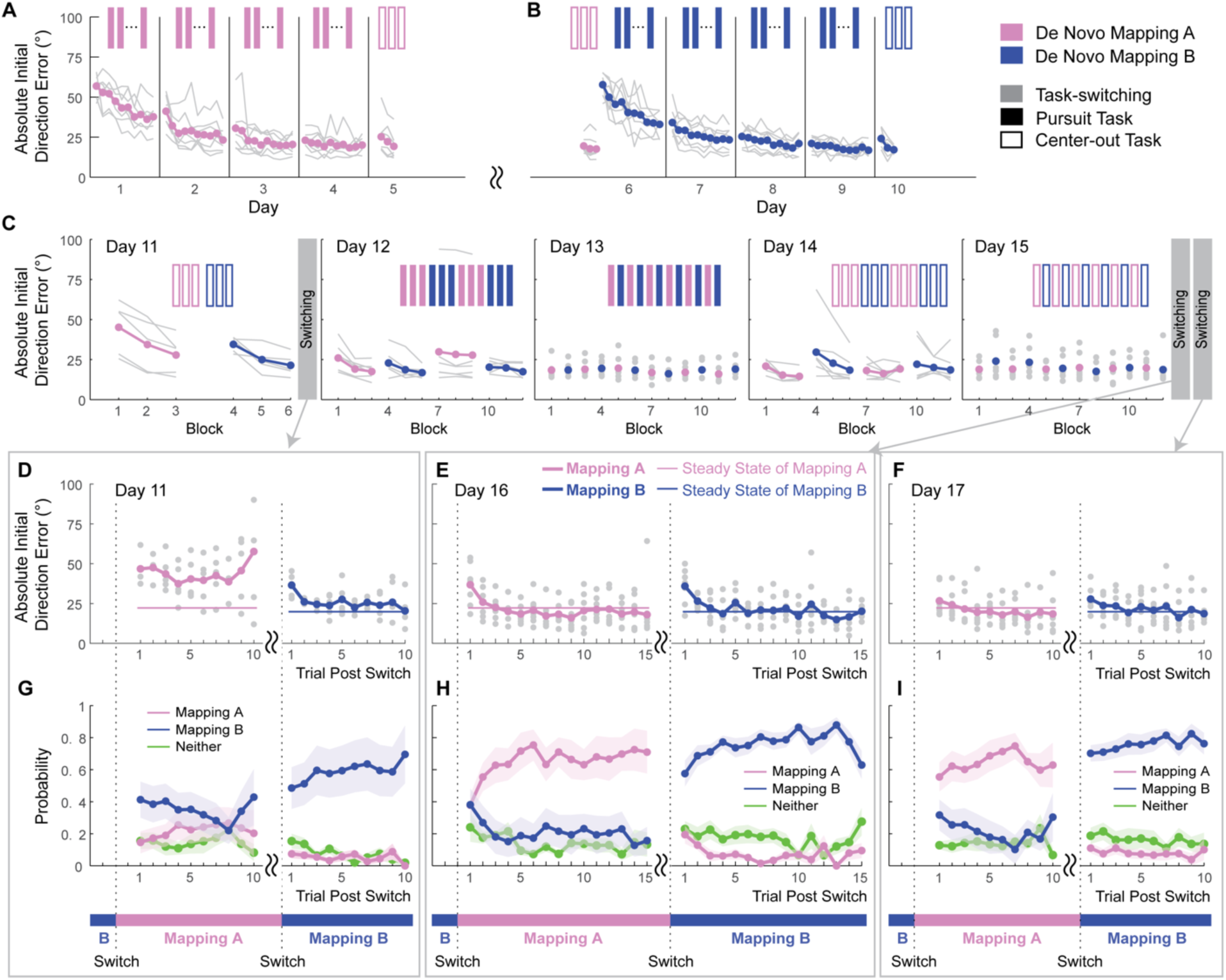
Learning and switching between two De Novo mappings. A,B) Performance improvement during initial training for each mapping (Days 1-10). C) Performance in subsequent training blocks that had a fixed mapping throughout, but with the mapping varying from block to block (Days 11-15). D-F) Performance (mean initial directional error) in blocks in which the mapping switched within-block (Days 11, 16, 17), plotted as a function of trial number relative to a switch. G-I) Probability of using each policy (established based on performance on Day 5 for Mapping A and on Day 10 for Mapping B) as a function of trial number relative to a switch. (D, G) Day 11 assessed switching performance immediately after learning both mappings, with the mapping switching every 2-10 trials. (E, H) Day 16 assessed switching performance after further training, with the mapping switching every 5-15 trials. (F, I) Day 17 repeated the same switch assessment as on Day 11, with the mapping switching every 2-10 trials. Thick magenta lines indicate average across participants for Mapping A, thick dark blue lines indicate those of the Mapping B. Gray dots/lines indicate data for individual participants. In the probability of choice, green thick lines indicate the probability neither Mapping A nor Mapping B were chosen. Shaded regions indicate ± s.e.m.

### Participants were unable to switch between two De Novo mappings shortly after initial learning, but could switch after more extensive practice

After they had successfully learned both Mapping A and Mapping B, we examined participants’ ability to switch between these two mappings (Fig. 4, Day 11). As in Experiment 1, we assessed this in a center-out version of the task, with the background screen color indicating the mapping to be used in the upcoming trial. This switched every 2-10 trials. Participants now exhibited substantially increased directional errors in the trials following a switch from Mapping B to Mapping A (Fig. 5D). Using the same approach as for Experiment 1, we used a maximum likelihood approach based on participants’ individual hand movement directions to estimate, on a trial-by-trial basis, the probability that participants used different policies post switch and determined the cause of errors: either persisting with the wrong policy or failing to correctly execute either policy. When switching from Mapping B to Mapping A, the probability of acting according to Mapping B remained high throughout the trials in which Mapping A was in use, while we found a relatively low probability that participants’ behavior was inconsistent with either the policy for Mapping A or the policy for Mapping B (Fig. 5G). In contrast, when switching from Mapping A to Mapping B, participants consistently acted appropriately for Mapping B (Fig. 5G). This asymmetry between mappings likely reflected a bias towards the most recently learned mapping.

Although participants struggled to switch between mappings shortly after they had learned both mappings, we wondered whether participants might be able to switch more readily if we allowed them some practice at switching between mappings. Over five days (Fig. 4, Days 12-16), participants practiced switching between Mapping A and Mapping B. This practice began with the mapping switching every three blocks (60 trials per block) while participants performed the pursuit task on Day 12, and then every block on Day 13. We then changed to a center-out task and switched the mapping every three blocks on Day 14, and every block on Day 15. On Day 16, participants practiced switching between mappings every 5-25 trials. Finally, on Day 17, we repeated the same task switching assessment as on Day 11, switching every 2-10 trials (Fig. 4, Day 17).

Participants showed smaller directional errors following a switch in the second task switching assessment on Day 17 in comparison to the switching assessment that was performed immediately after learning Mapping B on Day 11 (Fig. 5F). At a group level, when switching from Mapping A to Mapping B, participants showed increased initial direction errors in trials immediately following a switch, compared with steady state performance (steady state of the Mapping A (mean ± s.e.m): 19.87 ± 1.45 °, the first trials following switch: 27.80 ± 2.95 °, paired t-test, P = 0.0219, t(6) = 3.07). When switching from Mapping B to Mapping A, we didn’t see increased initial direction errors compared with steady state performance even from the first trials post switch (steady state of Mapping B (mean ± s.e.m): 22.22 ± 3.81 °, the first trials following switch: 26.68 ± 3.71 °, paired t-test, P = 0.3293, t(6) = 1.06). The behavior on Day 16, in which the mapping switched less frequently, was qualitatively similar to their behavior on Day 17 (Fig. 5EH).

As before, we examined participants’ hand movements to infer the cause of the increased errors following a switch. When switching from Mapping B to Mapping A, participants appropriately acted according to Mapping A with high probability on the very first post-switch trial. When switching from Mapping A to Mapping B, participants acted according to Mapping B with high probability (Fig. 5I). In both cases, the probability of not acting according to either policy remained approximately constant. As in Experiment 1, therefore, to the extent that there were switch costs, they were attributable to acting according to the wrong policy, not to errors in execution or implementation of the relevant policy.

## Discussion

Here, we tested whether participants could rapidly and repeatedly switch between a pre-existing baseline controller and a controller learned through *de novo* learning, when prompted by a color cue. We found that, in both switch directions (De Novo to Baseline and Baseline to De Novo), participants generally did not have much difficulty in switching their behavior according to these two mappings. Most participants did, however, exhibit some increased errors in the first 1-2 trials following a switch. By closely analyzing participants’ movements of both their hands, we determined that these errors were attributable to inappropriate persistence with the policy used before the switch of mapping, rather than imperfect execution of the policy associated with the new mapping.

We subsequently asked whether participants could learn multiple skills associated with different mappings, and switch between them. We found that participants were able to learn two De Novo mappings with minimal interference, but initially struggled to switch between these two mappings. However, after further training that included frequent switching between mappings, they became able to switch more easily. Similar to the first experiment, we found that the errors participants exhibited after a switch were mostly attributable to inappropriately persisting with the policy associated with the pre-switch mapping, rather than failing to correctly implement the appropriate policy for the post-switch mapping.

Task switching has been extensively studied in cognitive tasks (Jersild, 1927; Allport et al., 1994; Rogers and Monsell, 1995; Kiesel et al., 2010). These studies have primarily focused on transient increases in reaction time. However, this approach yields limited insight into the underlying reason for these switch costs since it is unclear why additional reaction time might be needed. In our experiments, we applied time pressure to participants through the threat of the target disappearing, with the goal of eliminating any transient increases in reaction time and instead revealing potential errors in action selection (Hardwick et al., 2019). This approach was largely successful; participants did not transiently increase their reaction time following a switch of the mapping and instead exhibited increased task errors. The continuous nature of the actions our task provided much richer information about the nature of participants’ errors compared to button-pressing tasks and allowed us to infer that errors were primarily due to inappropriate persistence with the wrong mapping. This interpretation is consistent with the idea that task switching in general is attributable to difficulty in overcoming the previously active task policy, rather than a difficulty in retrieving the one required after the task changes – often referred to as “task set inertia” (Allport et al., 1994; Wylie and Allport, 2000). Indeed, persistence of the components of pre-switch task set has been observed in cognitive tasks. In an fMRI experiment, reaction time costs of switching to one task were predicted by activation of the region where activated during another task, although different regions were activated during each task (Wylie et al., 2006; Yeung et al., 2006), which suggests that ‘task set inertia’ can at least partially explain switch costs. Our behavioral findings parallel these neuroimaging findings.

One caveat to our results is that most participants’ reaction times were overall elevated during the switching blocks compared to blocks without any switching. These increased reaction times did not, however, seem to reflect participants taking additional time in order to overcome the persistence of the pre-switch mapping. While one might expect that participants with longer reaction times would have been more successful in switching their behavior, we in fact found the opposite pattern: participants who had a greater increase in reaction time also tended to perform worse in trials immediately following a switch. These effects might therefore have reflected differences in overall task engagement across individuals.

Theories of cognitive task switching suggest that switch costs might reflect a reconfiguration of the prefrontal cortex circuitry to reflect the revised task demands (Kim et al., 2012; Worringer et al., 2019; Egner, 2023). This process may involve multiple cognitive processes, such as shifting attention between different stimulus attributes, retrieving goal states (what to do), and mapping goal states to actions (how to do it). When we switch between motor tasks, a qualitatively similar reconfiguration process may be necessary. In this setting, to change tasks, we have to reconfigure our motor system in an analogous way as for cognitive tasks – we may have to orient our attention to different sensory feedback, retrieve different goal states for the task, and retrieve a suitable policy for performing the task. Our experiments primarily emphasized the last of these processes. One major potential difference, however, between our motor skill task and typical cognitive tasks is that our motor skill tasks were likely highly automatized by participants (Yang et al., 2022), whereas rule-based cognitive tasks may be performed in a more deliberative manner, potentially making it more challenging to switch between tasks.

Our second experiment examined whether or not people could learn and switch between two different de novo skills. Participants found this very challenging at first, but became able to switch proficiently after some practice. The ability to switch between policies for different mappings might be affected by either extent of practice on the two tasks being performed or on the amount of practice one has had at switching between them. Learning new motor skills require multiple hours of practice to master and are known to undergo qualitative changes during practice. Performance of a new skill is thought to be highly deliberative and cognitive in early stages of learning, but become more automatic later in learning (Shiffrin and Dumais, 1981; Haith and Krakauer, 2018; Du et al., 2022). The cost of switching between different motor tasks might therefore depend on the mode of performance (i.e. deliberate versus automatic). Although we found that further practice allowed participants to better switching between mappings, it’s unclear whether this is attributable to a shift in the way each policy is implemented (i.e. deliberate versus automatic), greater experience in actually switching between policies, or something else.

The ease with which participants could switch between the De Novo mapping and the Baseline mapping stands in stark contrast to findings from adaptation, where it does not seem possible to revert to baseline control based on a color cue or other cue (Osu et al., 2004; Addou et al., 2011; Forano et al., 2021). This inability to switch is an inherent property of the implicit recalibration mechanisms involved in adaptation (Huberdeau et al., 2015; McDougle et al., 2016; Morehead et al., 2017; Krakauer et al., 2019). Indeed, this implicit recalibration mechanism is complemented by a further, explicit learning process, which is distinguished by the fact that it can be voluntarily disengaged through instruction. When countering very large perturbations, such as visuomotor rotations exceeding 90°, people do appear to be able to rapidly switch between two learned perturbations (Cunningham and Welch, 1994). In this instance, however, compensation is likely achieved through a simple re-aiming heuristic (Morehead et al., 2015; Wilterson and Taylor, 2021), rather than by making changes to the underlying controller. Therefore, although people can switch readily between opposing perturbations in, they likely do so by switching re-aiming strategies, rather than by switching to an entirely new controller. *De novo* learning tasks, by contrast, do not appear to depend on re-aiming strategies for learning (Yang et al., 2021) and, therefore, we believe that switching between *de novo* skills is quite different from switching between visuomotor rotations of different magnitudes.

Although not the primary focus of our study, our results demonstrate for the first time (to our knowledge) that participants can learn two distinct De Novo mappings with minimal interference – either retrograde (i.e. learning of Mapping B potentially disrupting the memory for Mapping A) or anterograde (i.e. prior learning of Mapping A making it easier or harder to subsequently learn Mapping B). Interference between motor memories has long been studied in the context of adaptation, with the goal of better understanding how multiple motor memories interact with one another (Brashers-Krug et al., 1996; Shadmehr and Brashers-Krug, 1997; Bock et al., 2001; Goedert and Willingham, 2002; Caithness et al., 2004; Krakauer et al., 2005; Krakauer and Shadmehr, 2006). Motor memories for adaptation are extremely weak, however. Adapted states are short-lived (Kitago et al., 2013), and so studies of long-term motor memory using adaptation paradigms had to focus on the fact that adaptation would be slightly faster the second time the same perturbation was experienced. Although much work continues to be generated that examines savings and interference in the context of adaptation paradigms, we suggest that *de novo* learning paradigms provide a more authentic approach to studying interference between motor memories since *de novo* learning much better parallels real-world skill learning which, unlike adaptation, remains largely stable with the passage of time. Although our experiment found no evidence for interference, it may be interesting to consider whether this might be different if episodes of practice for each mapping occur more closely in time, of if the two mappings are more closely aligned with one another.

## Materials and Methods

### Participants

A total of 25 participants took part in this study (25.77 ± 7.22 years old; 12 female). All participants took part in Experiment 1, and seven of these participants also took part in Experiment 2. All participants were right-handed and naive to the purposes of the study, had no known neurological disorder and provided written consent before participation. All procedures were approved by the Johns Hopkins University School of Medicine Institutional Review Board. All participants gave written informed consent and received $15 per hour for their participation.

### Experimental Setup

Participants sat on a chair in front of a glass-surfaced table with both arms resting on a plastic cuff mounted on an air sled which enabled frictionless planar movement of their arms across the surface of the table (Fig. 1A). Participants could not directly see their arms. Instead, they viewed visual cues (cursor and targets) which were displayed in the plane of the hand through a mirror positioned horizontally above their arms. The position of both the left and right hands was tracked at 130 Hz using a magnetic tracking device (Flock of Birds; Ascension Technologies).

### Experiment 1

#### Training for the De Novo mapping

In Experiment 1, 25 participants completed five sessions, each on separate days. In the first four days, participants learned to control the cursor under the De Novo mapping by making a series of point-to-point movements around the workspace (Pursuit Task, Fig. 1B). On each block, participants were required to move their hands to guide an on-screen cursor (5 mm diameter) to a fixed start circle (10 mm diameter) appearing in the center of the workspace. Once participants held the cursor stationary inside the start circle, a target appeared at a location 12 cm from the start position. After participants reaching this target, a new target appeared at different location. The distance between targets was always 12 cm, but the direction was determined pesudorandomly within a 20 cm x 20 cm workspace centered on the start location. We used two mappings (Fig. 1C): the Baseline mapping, in which the cursor appeared at the average location of the two hands and was therefore intuitive to use, and the De Novo mapping which involved a non-intuitive mapping between hands and cursor that had to be learned. In the De Novo mapping, forward-backward movement of the left hand controlled left-right movement of the cursor, and left-right movement of the right hand controlled forward-backward movement of the cursor (Haith et al., 2022). Both mappings were explicitly explained to participants at the start of the experiment. On the first day, participants were not given any specific instruction when they practiced the De Novo mapping, but from the second day, they were asked to move the cursor in a straight line between targets as best as they could and to maintain their peak movement speed within a particular range (0.39 – 0.52 m/s). Feedback about movement speed was given by changing the color of the target at the end of the movement. If the peak speed was within the required range, the target changed color from gray to yellow and a pleasant sound played. If the movement speed was too fast, the target changed color from gray to magenta and, if the movement was too slow, it changed color to blue. Participants performed the task in a series of blocks, each of which consisted of 60 point-to-point movements. On the first day, participants performed three blocks of the pursuit task under the Baseline mapping to become familiar with the task and experimental setup, and then practiced the De Novo mapping for ten blocks. From the second to fourth day, participants performed one block with the Baseline mapping at the start of the session, and then continued practicing the De Novo mapping for ten blocks (Fig. 1D, Training).

#### Assessment of switching between mappings

On the fifth day, we examined participants’ ability to switch between the learned De Novo mapping and the Baseline mapping (Fig. 1D, Testing). To facilitate switching between mappings and to ensure that initial conditions could be easily matched across mappings, participants made center-out movements from a fixed starting position in each trial (Center-out Task, Fig. 1B). Participants returned their hands to the same starting positions after each movement. We positioned air nozzles above the starting position for each hand to help guide participants back to the appropriate starting posture. If both left and right hand were within 50 mm from the starting position for each hand, a trial started. During the center-out task, the background color of the screen turned either blue or magenta to indicate which mapping should be used in the upcoming trial, visible 500 ms before the target location was presented. The background color-mapping association did not randomize across participants, light blue and magenta indicated the Baseline and the De Novo mapping, respectively.

To ensure participants reacted rapidly to the appearance of each target, we applied time pressure through the threat of the target disappearing. After the target appeared, which served as the cue to begin moving, the target disappeared at a random, exponentially-distributed time (0.5 – 2 s) within each trial, after which they were unable to successfully acquire the target. Participants first practiced this center-out task under either the Baseline mapping (1 block of 60 trials) or the De Novo mapping (3 blocks of 60 trials).

We then assessed participants’ ability to switch between the Baseline mapping and the De Novo mapping. On each trial, the cursor was controlled either through the newly learned De Novo mapping, or through the Baseline mapping and the mapping for the upcoming trial was indicated by the color-cue (Fig. 1E). The mapping switched randomly every 2-10 trials. Participants completed 10 blocks. Each block consisted of more than 63 trials, during which time the mapping switched 5 times in each direction (i.e., 5 times from Baseline to De Novo and 5 times from De Novo to Baseline).

### Experiment 2

In Experiment 1 we examined participants’ ability to switch between a newly learned mapping (De Novo mapping) and a highly overlearned mapping (Baseline mapping). In Experiment 2, we tested whether participants could learn two De Novo mappings and be able to switch between them.

#### Practice for the second De Novo mapping

After Experiment 1, seven of the 25 participants from Experiment 1 joined Experiment 2. In addition to the De Novo mapping that participants learned in Experiment 1 (which we will refer to as Mapping A), participants practiced a second De Novo mapping (Mapping B, Fig. 4). In this mapping, the contingencies between hand movement and cursor movement were transposed compared to Mapping A; forward-backward movement of the right hand controlled left-right movement of the cursor, and left-right movement of the left hand controlled forward-backward movement of the cursor. The structure of Mapping B was explicitly explained to participants at the start of Day 6. Days 6-10 of Experiment 2 were structured exactly the same as Days 1-5 in Experiment 1 (Fig. 1D, Fig. 4, Days 6-10).

#### Assessment of switching between two De Novo mappings

Before testing whether participants could switch between Mapping A and Mapping B, we first confirmed that participants could still successfully perform under Mapping A. After a warmup block for each mapping under by pursuit task, participants performed center-out task under either Mapping A or Mapping B (3 blocks of 60 trials for each mapping, Fig. 4, Day 11).

We then assessed whether participants could switch between Mapping A and Mapping B (Fig. 4, Day 11), following exactly the same approach as in Experiment 1. Since we found that participants struggled to switch immediately after learning both mappings, we gave participants an opportunity to practice switching between mappings. Participants practiced switching between Mapping A and Mapping B over five days, with the frequency of switching steadily increasing over days (Fig. 4, Days 12-16). Participants switched between mappings every three blocks with pursuit task on Day 12, every block with pursuit task on Day 13, every three blocks with center-out task on Day 14, every block with center-out task on Day 15, every 5-25 trials with center-out task (10 blocks of at least 63 trials) on Day 16. Participants performed 6 blocks of 60 trials for each mapping in total on Days 12-15. Then we repeated the same switching assessment from Day 11 which matched that in Experiment 1 (Fig. 4, Day 17). Two of seven participants did not complete the initial switching assessment on Day 11 and instead directly started switching training (Days 12-16).

### Data analysis

Raw data of both hands position were smoothed using a Savitzky–Golay filter to eliminate high-frequency measurement noise, differentiated to obtain movement velocity and then smoothed again. Movement onset was determined based on the first time that the tangential velocity of the hand exceeded 0.026 m/s. Then the delays in our system (measured to be 100 ms) were subtracted from this time to obtain an estimate of the true time of movement initiation relative to the target appearing on the screen. The participant’s reaction time in each trial was determined as the delay between the time of stimulus presentation and the time of movement initiation. Initial movement direction in each trial was defined based on the direction of the velocity vector of the hand 100 ms after movement onset.

To observe switch costs in initial movement directional errors, initial direction errors during center-out task were used as steady state performance for each mapping (Day 5 for the Baseline mapping and the De Novo Mapping A, Day 10 for the De Novo Mapping B). We compared the initial direction errors between these steady state performance and trials post switch.

We developed a probabilistic approach to estimate which policy participants attempted to use on each trial during the switching assessment, based on the initial movement direction of the right and left hands. We assumed that the distribution of left and right initial hand directions given the target direction comprised a mixture of three different possible policies: two associated with each mapping (Mapping A and Mapping B), plus a third, random policy that did not conform to either mapping:

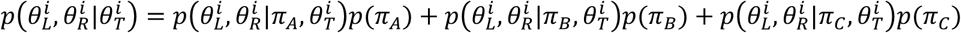

Here 𝜋 indicates the policy participants used (𝜋*_A_*: 𝐵𝑎𝑠𝑒𝑙𝑖𝑛𝑒, *𝜋_B_*: 𝐷𝑒 𝑁𝑜𝑣𝑜, 𝜋*_C_*: 𝑂𝑡ℎ𝑒𝑟 (uniform distribution) in Experiment 1, 𝜋*_A_*: 𝑀𝑎𝑝𝑝𝑖𝑛𝑔 𝐴, 𝜋*_B_*: 𝑀𝑎𝑝𝑝𝑖𝑛𝑔 𝐵, 𝜋*_C_*: 𝑂𝑡ℎ𝑒𝑟 in Experiment 2), 𝜃*_R_* and 𝜃*_L_* are the initial directions of the right and left hands, and 𝜃*_T_* is the direction of the target. 𝑝(𝜃*_L_*, 𝜃*_R_*|𝜋, 𝜃*_T_*) is the distribution of the left and right hand movements under each policy.

We estimated the distributions associated with Mapping A and Mapping B separately for each participant via kernel density estimation based on center-out task blocks at Day 5 for Mapping A and Baseline and those at Day 10 for Mapping B in Experiment 2. For simplicity, we assumed that the left and right hand movements were independent given the policy and target direction (i.e. 𝑝(𝜃*_L_*, 𝜃*_R_*|𝜋, 𝜃*_T_*) = 𝑝(𝜃*_L_*|𝜋, 𝜃*_T_*)𝑝(𝜃*_R_*|𝜋, 𝜃*_T_*)). The overall probability of a participants behavior across multiples trials was therefore given by:

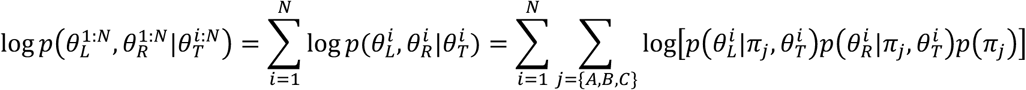

We used this model to estimate the probabilities of each policy 𝑝(𝜋_𝐴_), 𝑝(𝜋_𝐵_) and 𝑝(𝜋_𝐶_) by maximum likelihood, constraining 𝑝(𝜋_𝐴_), 𝑝(𝜋_𝐵_), 𝑝(𝜋_𝐶_) to be between 0 and 1 and requiring that 𝑝(𝜋*_A_*) + 𝑝(𝜋*_B_*) + 𝑝(𝜋*_C_*) = 1. We separately estimated these policy probabilities at different numbers of trials post-switch, pooling data from all trials at a given trial number after a switch (1st through 10th trials post switching for Experiment 1 and at 1st through 15th trials post switching for Experiment 2).

Our approach is illustrated through example trials in Figure 3A, which shows: (1) trial *i* is located on the distribution of the De Novo mapping, and is therefore highly likely to have been generated by the De Novo policy. (2) trial *j* is located on the distribution of the Baseline behavior and is therefore likely to have been generated under the Baseline policy. (3) trial *k* does not overlap with either the De Novo or Baseline distributions. This trial is better explained by a third, uniform distribution. This analysis therefore allowed us to quantify the probability of using different policies, and therefore determine the nature of errors. Analysis was performed using MATLAB R2022a (MathWorks).

### Statistics

Unless otherwise specified, two-tailed paired t-tests were used to assess differences in behavior across conditions (for example, the Baseline mapping vs. the De Novo mapping), and two sample t-tests were used to assess differences between steady state behavior and behavior post switching (for example, reaction time at center-out task vs. reaction time at trials post switch). All statistical tests were conducted in MATLAB R2022a.

## Acknowledgments

This work was supported by a grant from Meta Reality Labs and the Sheikh Khalifa Stroke Institute.

